# Analysis of Wheat Spike Morphological Traits Using 2D Imaging

**DOI:** 10.1101/2025.06.06.658159

**Authors:** Fujun Sun, Shusong Zheng, Zongyang Li, Qi Gao, Ni Jiang

## Abstract

The morphological structure of wheat spikes plays a central role in wheat yield. Wheat spike morphology, closely associated with crop yield, has attracted considerable attention in the fields of genetics and breeding. However, traditional measurement methods can only measure simple traits, and precise phenotypes remain difficult to obtain, constraining the study and improvement of complex spike-related traits. This study utilized deep learning technologies to develop a pipeline, called SpikePheno, for the acquisition of precise wheat spike phenotypes. Our pipeline demonstrated high accuracy in spike segmentation, achieving a mean Intersection over Union (mIoU) of 0.948. Additionally, our method accurately identified spikelet counts, achieving an R^2^ of 0.9923. Using experimental data of 221 wheat cultivars from various regions of China grown in Zhao County, Hebei Province, our pipeline extracted 45 different phenotypes and studied their correlations with thousand grain weight (TGW) and spike yield. Our findings indicate that precise measurement of spike area, spikelet area, and other phenotypic traits enables a clearer understanding of the correlation between spike morphology and wheat yield. Through hierarchical clustering based on spike morphology, we categorized wheat spikes into six classes and identified phenotypic differences between these classes and their impact on TGW and yield. Furthermore, this study revealed phenotypic differences between wheat cultivars from different geographical regions and over different decades, with an increase in large-spike cultivars over time, especially in southern China. This research may help breeders understand the relationship between wheat spike morphology and yield, providing an important basis for future wheat breeding efforts.

## 1 Introduction

Wheat (*Triticum aestivum* L.) plays a major role in the global food supply. As the global population grows, increasing wheat yield has become a key priority in agricultural research to ensure food security. Among the critical determinants of wheat production, spike number per area (SNA), grain number per spike (GNS), and thousand grain weight (TGW) are essential. The major yield components in wheat often exhibit compensatory effects. Coupled with the polyploid nature of wheat, these effects complicate the genetic and physiological studies of these traits Brinton and Uauy (1). The complex competitive interactions among SNA, GNS, and TGW make achieving a singular yield enhancement target challenging. Given the central role of spike structure in the formation of wheat yield, studying it in depth is not only essential for understanding yield potential but also a crucial foundation for breeding optimization.

In the morphological structure of wheat spikes, the spikelet number per spike (SNS) is a critical determinant of GNS [2]. Each spikelet typically contains 5-7 florets, each with the potential to develop into a mature grain. Researches have shown that by increasing SNS or improving the fertility of florets, one can optimize inflorescence architecture, thereby directly enhancing GNS and thus overall crop yield [3]. Additionally, the optimization of spikelet structure can enhance photosynthesis and water utilization efficiency, further increasing wheat productivity [4]. Studies have found that the removal of spikelets typically results in a slight increase in the number of grains in the remaining spikelets and a significant increase in grain weight [5]. Thus, the design of spike architecture requires careful regulation to achieve optimal yield and quality [6]. In breeding practices, molecular marker-assisted selection and gene editing technologies allow researchers to precisely manipulate genes affecting spikelet development, thereby optimizing spike architecture for enhanced yield potential[7].

In addition to influencing grain yield, the structure of wheat spikes also affects the plant’s resistance to diseases and pests [8]. Compact spikes, characterized by smaller spikelet gaps, are more susceptible to certain fungal diseases due to their specific microclimate conditions, such as humidity and temperature. After rainfall or irrigation, compact spikes restrict air circulation due to the close packing of spikelets, making it difficult for moisture to evaporate or drain away quickly. This environment fosters the growth and proliferation of pathogenic fungi [9, 10]. Moreover, the spike plays a crucial role in wheat’s ability to withstand abiotic stress. Under high temperature conditions, long-spike wheats exhibit better heat tolerance because the spike photosynthesis continues to supply essential assimilates to the grains, even when the leaves senesce due to heat, thereby supporting grain growth and development and ensuring yield stability [11]. Therefore, a well-structured spike is crucial for achieving high yield.

Traditional spike phenotyping relies on manual methods. Researchers use rulers to measure spike length (SL) and manually count SNS and GNS. This approach is not only time-consuming and labor-intensive but also prone to human error, posing significant challenges to the accuracy and efficiency of spike phenotyping. Imaging technologies have greatly enhanced the ability to capture wheat spike traits efficiently. These technologies allow for the automatic extraction of traits from spike images. Recent advancements in deep learning models have further improved the accuracy of spike segmentation and trait extraction. Batin, Islam (12) applied the Cascade Mask R-CNN for segmenting and counting wheat spikes from field imagery with bounding box and mask mean average precision (mAP) scores of 0.9303 and 0.9416, respectively. [13] presented an improved Fully Convolutional Network (FCN) to segment spike regions and applied a density estimation method to count wheat spikelets from infield images. The symmetric mean absolute percentage error (SMAPE) for SNS estimation ranged from 36.52 to 97.13 across different growth stage in the prediction set. Despite advancements in phenotyping spikes in field environments, accurately phenotyping spike characteristics such as SNS, grain size, and fertility remains challenging due to complex backgrounds and spikelet occlusion. For detailed and accurate spike trait measurements, the spike phenotyping from spike images is often chosen. For example, Qiu, He (14) used the Faster Region-based Convolutional Network (Faster R-CNN) to detect and count wheat spikelets in spike side view images, with root mean squared errors (RMSE) ranged from 0.54 to 0.77. However, it could not segment individual spikelets or extract specific spikelet traits, limiting its applicability for detailed wheat spike analysis. Niu, Liang (15) applied the YOLOv8-seg to segment spikelets from wheat frontal view images, and classified the number of grains in each spikelet with Support Vector Machine (SVM) model, achieving a mean absolute error (MAE) with 1.04 for GNS in spike images. However, the deep learning models in the above studies were trained on images from only a few cultivars, which limits their generalizability to diverse cultivars with varying spike traits. These models may be overfit to the specific characteristics of the training cultivars and fail to accurately distinguish between fertile and infertile spikelets when applied to different varieties. The X-ray Computed Tomography (CT) imaging offers precise and detailed 3D visualization of wheat spike architecture [16, 17]. While it provides excellent capability in measuring wheat spikes, spikelet and grain morphological traits, the high cost and low throughput of CT imaging limit its widespread use in large-scale phenotyping.

The main objective of this study was to develop a rapid and low-cost phenotyping method to analyze the complex structure of wheat spikes. To achieve this, we proposed a pipeline called SpikePheno. A semantic segmentation model was trained to segment wheat spike regions. Overall spike morphological characteristics including shape, size and area were obtained. Additionally, an instance segmentation model was trained to segment individual spikelets from spike regions, enabling the extraction of the number, morphology, and distribution of the spikelets. Our method is expected to provide a more efficient and accurate solution for wheat spike phenotyping and contribute to the development of high-yielding and resilient wheat cultivars.

## 2 Materials and Methods

### 2.1 Experimental materials and image acquisition

In the current study, 221 common wheat (*Triticum aestivum* L.) accessions were cultivated, comprising 6 landraces, 209 cultivars originating from China, and 6 introduced cultivars. These accessions were grown in Zhao County, Shijiazhuang City, Hebei Province, China (38°05′N, 114°52′E) during the 2022-2023 wheat growing season. The seeds were sown on October 9-10, 2022. The experimental fields were laid out according to an alpha-lattice incomplete block design with two replications, following the methodology described by [18]. Each plot was hand-planted with five rows of wheat, with each plot designed to be 6 meters in length, a 25 cm spacing between rows, and a 5 cm spacing between plants within each row. The field was managed according to local practices. To minimize interaction between adjacent plots, a 50 cm buffer zone was established. Detailed source information for the wheat cultivars is delineated in Table S2.

We set up an imaging station for spike phenotyping, equipped with an Allied Vision 1800 U-1620C camera mounted on a tripod. Two supplementary lights were used to provide optimal illumination. A black PVC plastic sheet was used as the background to enhance contrast, and a circular chip (2.5 cm in diameter) was placed on the sheet as a size marker. During imaging, the spikes were placed on their side. A total of 2198 images were collected. The size of each image was 4112×3008 pixels.

### 2.2 Spike segmentation model

In this study, we trained a segmentation model to segment the wheat spike and stem from the background based on the UNet architecture (Fig. 1A) [19]. To enhance model performance, the backbone network was replaced with ResNet50 [20]. It introduced the residual structure using shortcut connections to address the problem of vanishing and exploding gradients as the network depth increases. The residual block used a bottleneck design, where each block consisted of three convolutional layers: a 1×1 convolution to reduce dimensions, a 3×3 convolution for feature extraction, and another 1×1 convolution to restore dimensions (Fig. 1B). Moreover, at the beginning of each stage, the first bottleneck block adjusted the dimensions using a 1×1 convolution layer when the input and output feature channels are not the same.

**Fig. 1.**
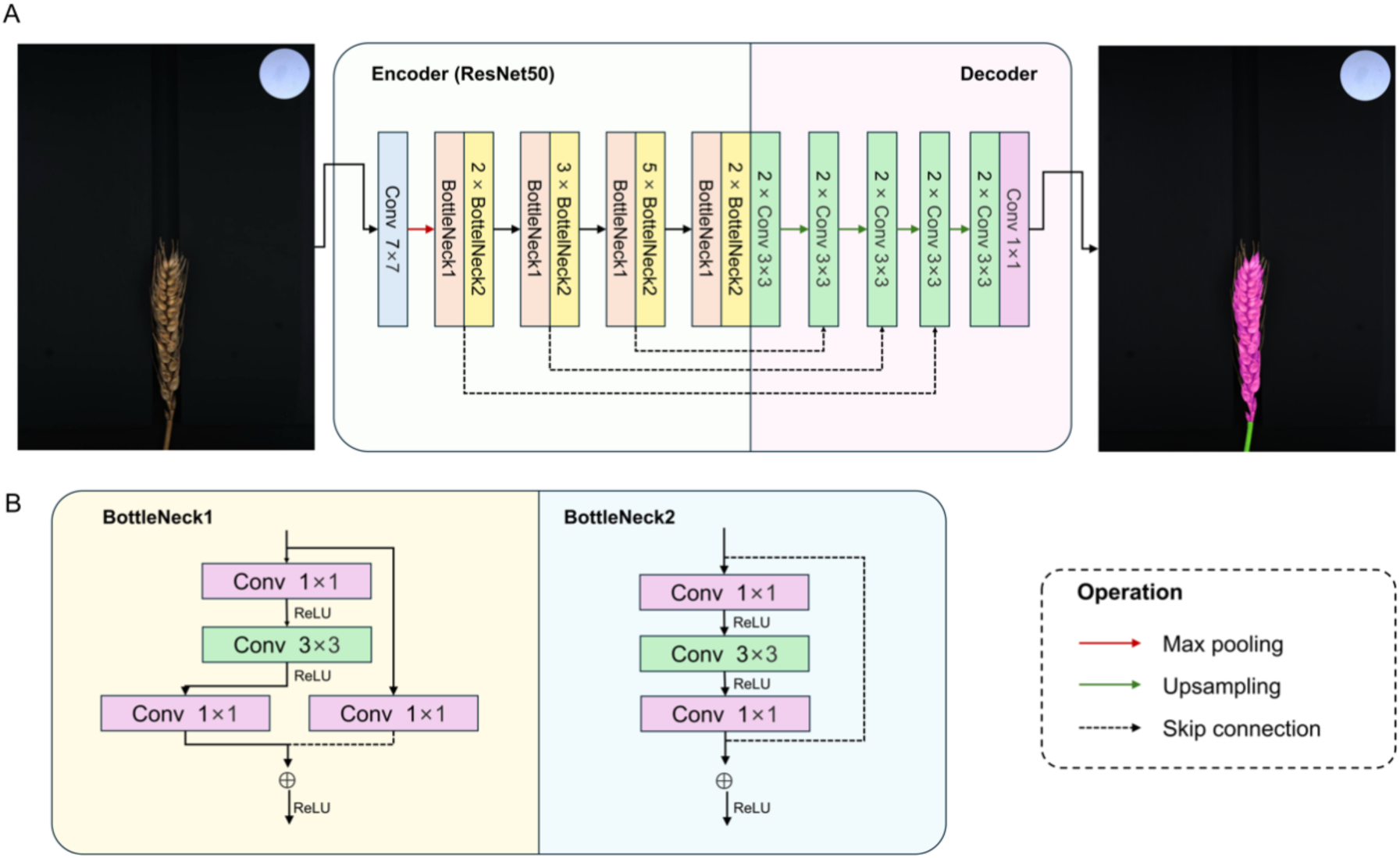
Segmentation of wheat spike and stem from background using ResNet50-UNet. (A) Architecture of ResNet50-UNet. (B) BottleNecks of ResNet50.

During training, images were augmented through random scaling, color perturbation, and flipping. We defined the loss function *L* as a combination of a weighted Focal Loss function and Dice Loss:

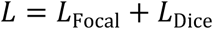

The weighted Focal Loss is defined as follows:

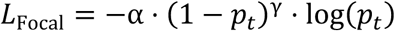

where *p*_i_ is the model’s predicted probability of a sample belonging to the positive class, α is the sample weight set to 0.5, and γ is the tuning parameter set to 2. The Dice Loss is defined as:

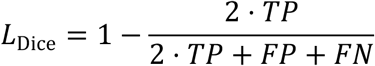

where *TP* represents the correctly predicted positive pixels, *FP* denotes the incorrectly predicted negative pixels as positive, and *FN* means the incorrectly predicted positive pixels as negative.

We used a dataset consisting of 80 images of wheat spikes for training and evaluating the spike segmentation model. The dataset was divided into a training set (50 images), a validation set (10 images), and a test set (20 images). To ensure the model’s generalization capability, the test set was specifically designed to include half of the images from wheat cultivars that were included in the training and validation sets (hereafter referred to as seen cultivars) and the other half from wheat cultivars that were not included in these sets (hereafter referred to as unseen cultivars). To preserve important features and optimize GPU memory usage, the images were zero-padded and resized to 1024×1024 pixels. We employed the Adam optimizer for training. To enhance training stability, we implemented a learning rate adjustment strategy that included a warm-up phase followed by cosine annealing [21]. The initial learning rate was set at 1×10^-4^, and the minimum learning rate was set at 4×10^-6^. The model was trained for 850 epochs.

### 2.3 Spikelet segmentation model

YOLOv8-seg, an instance segmentation extension of the YOLOv8 model, retains the advantages of the YOLO series in object detection and is specifically optimized for instance segmentation tasks [22]. In this study, we employed YOLOv8x-seg to train a spikelet segmentation model, which was designed to segment individual spikelets from spike images and classify them as fertile or infertile (Fig. 2). The spikelet segmentation model used an improved backbone structure that introduced the CSP Bottleneck with 2 Convolutions (C2f) module for multi-scale feature fusion, enhancing feature extraction capabilities. In the Neck part, it adopted a PAN-FAN structure, which combined the Path Aggregation Network (PAN) [23] and Feature Aggregation Network (FAN) [24] to perform multi-scale feature fusion, improving the model’s ability to detect objects at different scales. On top of the detection head, a segmentation head inspired by YOLACT [25] was added, which generated feature maps for mask prediction.

**Fig. 2.**
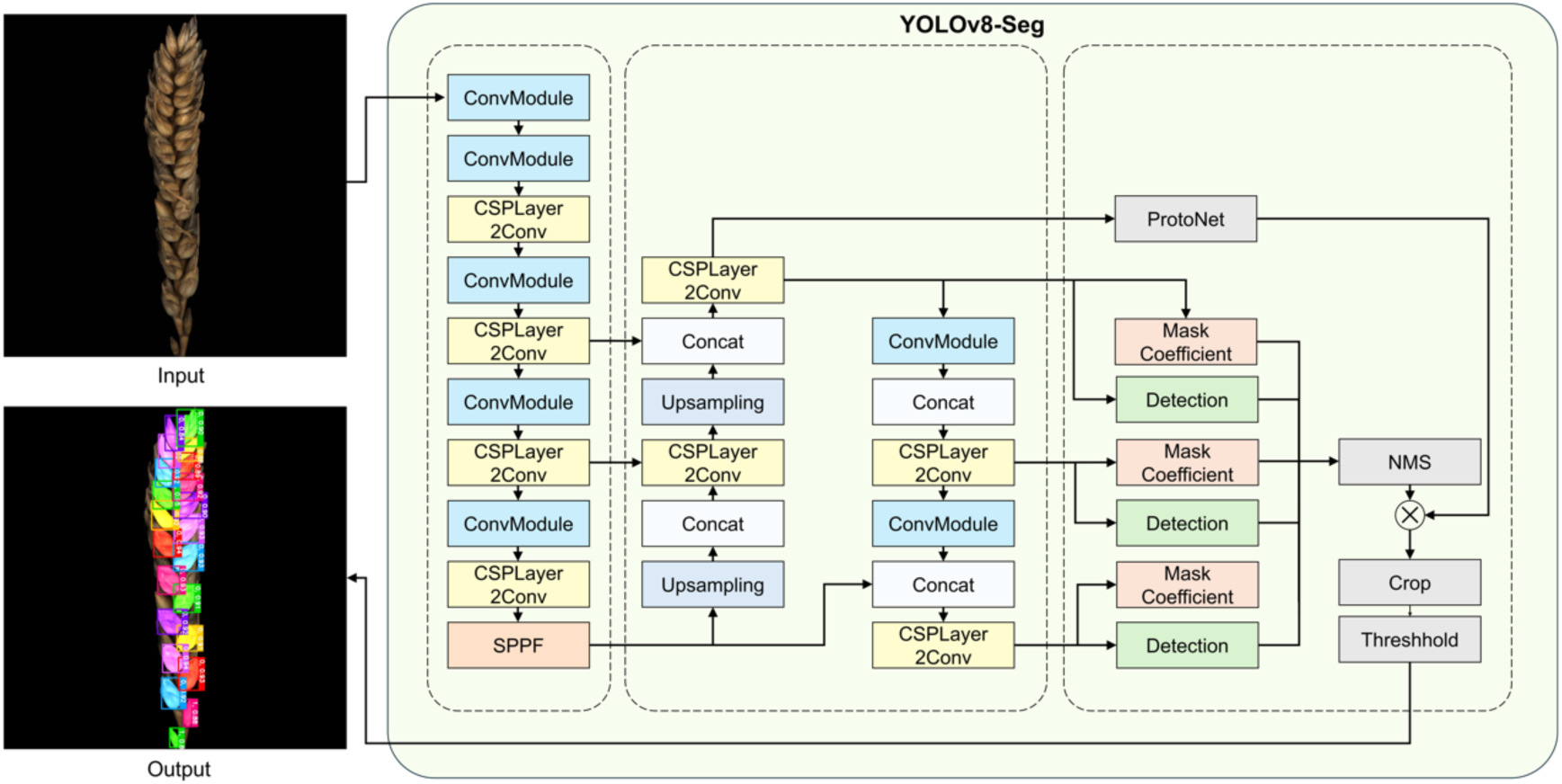
Segmentation of spikelets from the wheat spike.

Using the output mask images generated by the spike segmentation model, the spike images were cropped. The cropped images were then zero-padded to a square size and resized to 960×960 pixels before being input into the spikelet segmentation model. We conducted experiments using a dataset consisting of 330 images of wheat spikes for the spikelet instance segmentation model. The dataset was divided into a training set (240 images), a validation set (30 images), and a test set (60 images). The test set was specifically designed to include half of the images from wheat cultivars seen by the model and the other half from unseen cultivars. During the training process, images were augmented through translation transformation, color distortion, random rotation, and vertical flipping to enhance model performance. The model was trained using Adam optimizer with a learning rate 1×10^-3^. It was trained for 600 epochs, and the epoch with the highest validation performance was kept.

### 2.4 Morphological traits extraction

The overall spike traits, as well as traits for individual spikelets, were derived from segmented image masks. The segmented spike mask was used to compute spike area, perimeter, length, width, and roundness, and the individual masks of spikelets were used to derive estimates of spikelet number, width, angles, and density.

For each spike images, the spike and stem were automatically segmented into regions using the spike segmentation model (Fig. 3A). First, we computed the skeleton of wheat spike using Zhang-Suen thinning algorithm [26] based on the extracted spike mask. Due to the natural irregularities in the shape of wheat spikes, the extracted skeleton might have redundant branches or noise. To address this, we recorded the skeleton coordinates and identified the points closest to the top and base of the wheat spike region (Fig. 3C). The skeleton coordinates were then processed using the Depth-First Search (DFS) traversal algorithm to form a path from the top point to the base point along the skeleton. Additionally, to reduce noise, this path was smoothed using the Savitzky-Golay filter [27] with linear fitting and a window size set to half of the total number of the path points. To minimize the edge effects of the filter, which can result in variations in the length of the path at both ends, we implemented an optimization strategy by extending the pixels at both ends of the path. The slope information from the 10 pixels at each end was used for extension. This approach allowed us to obtain an optimized skeleton representing the main axis of the spike. Based on the optimized skeleton, SL was computed as:

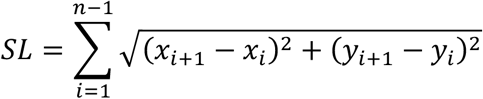

**Fig. 3.**
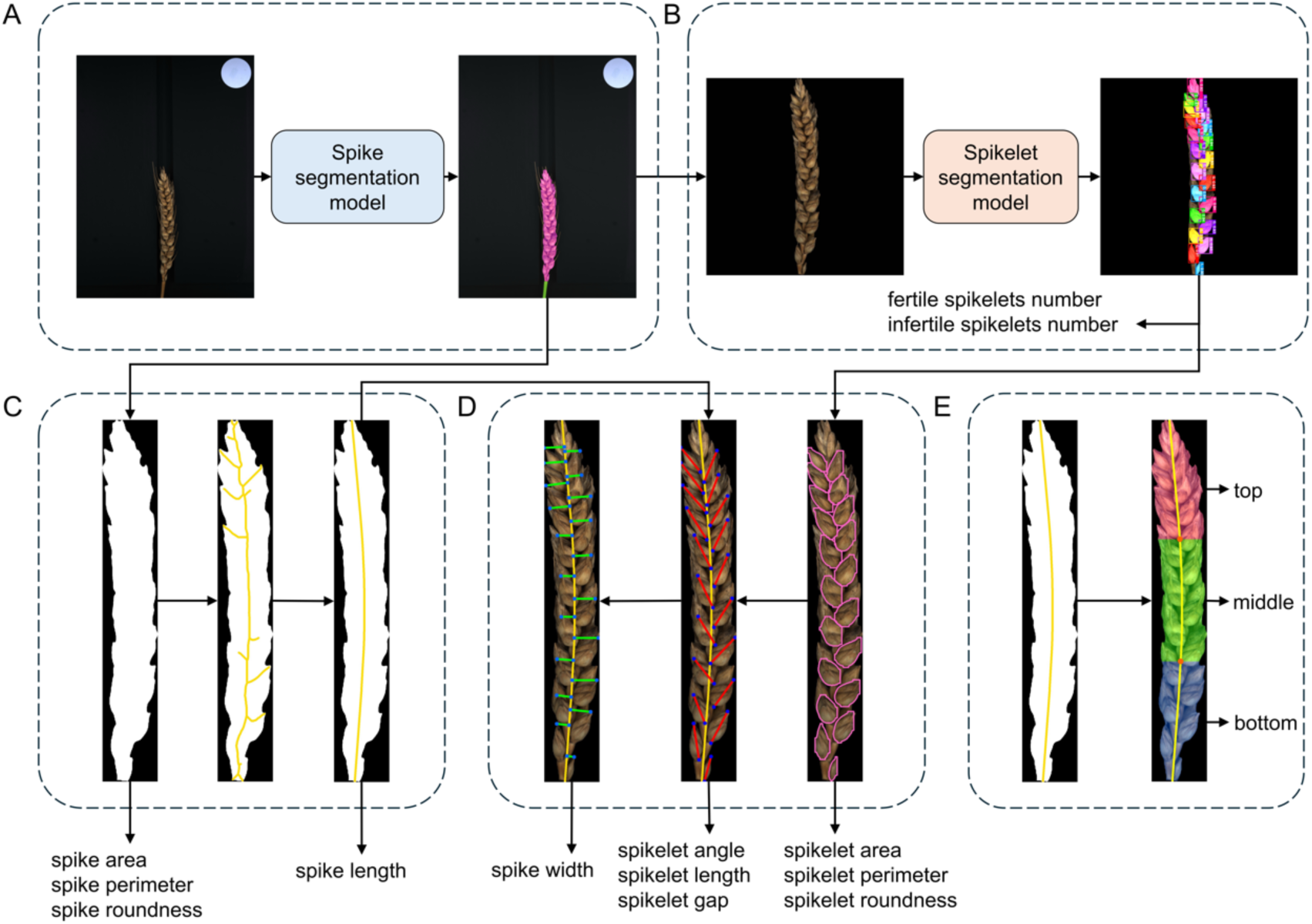
Workflow of spike and spikelets phenotyping (A) Segmentation of the wheat spike from the background. (B) Segmentation of wheat spikelets from the wheat spike. (C) Spike phenotyping using the main axis and the skeleton of the spike. (D) Spike and spikelet phenotyping based on the main axis of the spike and segmented spikelet masks. (E) Division of the wheat spike into three uniform segments by the main axis of the spike.

where *n* is the total number of points on the optimized skeleton, and (*xi*, *yi*) and (*xi+1, yi+1*) represent the coordinates of two consecutive points on the skeleton. Since the optimized skeleton reflected the natural curvature of spike, it enabled accurate length measurements, even when the spikes bend or was not straight.

The algorithms arclength and contourArea provided by OpenCV were used to compute spike perimeter (SP) and area (SA) respectively. The SA was approximated by the number of non-zero pixels in the segmented spike region. The SP was approximated by the boundary of segmented spike region. A roundness (SR) index was computed to describe the overall shape of the spike:

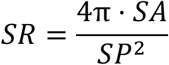

Based on the instance segmentation model for spikelets, we identified and counted both fertile and infertile spikelets (Fig. 3B). For each spikelet, similar to the spike, we computed the area, perimeter, and roundness using the segmented masks of the individual spikelets.

Principal Component Analysis (PCA) was applied to determine the main orientation of a spikelet based on its pixel coordinates (Fig. 3D). The pixel coordinates of the segmented spikelet mask were treated as data points, and the covariance matrix of these points was computed. The eigenvector corresponding to the largest eigenvalue (the first principal component), represented the direction of the greatest variance, which defined the major axis of the spikelet.

The spikelet length was estimated by projecting all pixel coordinates of the segmented spikelet onto its major axis (Fig. 3D). The two points with the maximum Euclidean distance along this axis were identified as the endpoints of the spikelet, and the distance between them was calculated as the spikelet length.

To measure the spikelet angle, the intersection point between the major axis of the spikelet and the main axis of the spike was determined by solving the equations of their respective lines in the image coordinate space (Fig. 3D). Since the main axis of the spike was optimized and smoothed, the local direction of the spike axis at the intersection point was interpolated. The spikelet angle was then computed as the angle *θ* between the major axis vector *vs* of the spikelet and the local direction vector *vv* of the spike:

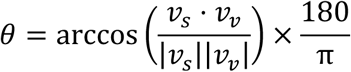

Due to the irregularity of the topmost spikelet of spike, we excluded it from phenotypic calculations, including length, area, perimeter, angle and roundness, as well as spike width (SW).

The spikelet gap was defined as the distance between the bases of two adjacent spikelets along the main axis of the spike (Fig. 3D). This gap was computed by identifying the base points of adjacent spikelets and calculating the Euclidean distance between them along the spike axis.

Additionally, we utilized the positions of fertile spikelets to measure SW (Fig. 3D). Since the shape of infertile spikelets could affect the overall structural characteristics of the spike, they were excluded from the SW measurement. For each fertile spikelet, the top endpoint of its major axis was identified, and the shortest distance from this point to the main axis was calculated. This distance was then doubled to estimate the local width of the spike at that position. The average SW across different regions of the spike was obtained by averaging the calculated widths from all fertile spikelets.

To further investigate the spatial distribution of spikelet traits along the spike, the spike was divided into three segments based on the extracted SL (Fig. 3E). Specifically, the length of the optimized skeleton representing the main axis of the spike was uniformly divided into top, middle, and bottom segments. Spikelet traits were quantified for each segment, which contained spikelets whose bases fell within the corresponding length range along the main axis. Detailed information on the spike and spikelets traits is provided in Table S1.

### 2.5 Evaluation metrics

To evaluate the performance of the spike segmentation model, we used mean Pixel Accuracy (mPA) and mean Intersection over Union (mIoU) as key metrics. The mPA measures the accuracy of pixel-wise classification by computing the average proportion of correctly classified pixels across all classes, while the IoU quantifies the spatial overlap between the predicted segmentation mask and the ground truth mask, indicating their alignment. These evaluation metrics are computed as follows:

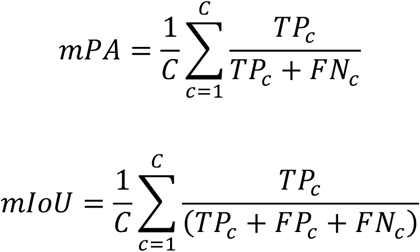

where *C* is the number of the classes, *TP* (True Positive) is the number of positive pixels classified accurately, *FP* (False Positive) is the number of actual negative pixels classified as positive, and *FN* (False Negative) is the number of actual positive pixels classified as negative.

The performance of spikelet instance segmentation model was evaluated using mean Average Precision with 0.5 IoU threshold for box detection (mAP50box) and mean Average Precision with 0.5 IoU threshold for segmentation (mAP50seg) as key metrics.

The mAP50box and mAP50seg at an IoU threshold of 0.5 are computed as follows:

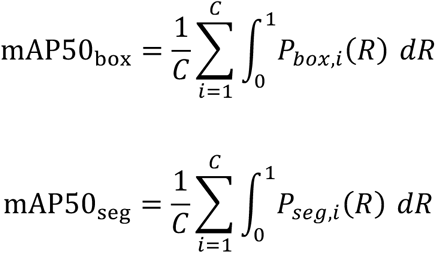

where *Pbox,i*(*R*) and *Pseg,i*(*R*) represent the precision-recall curves for bounding box detection and segmentation of class *i*, respectively, and *C* is the total number of classes. The mAP50box measures the localization performance based on the overlap between predicted and ground truth bounding boxes, while mAP50seg quantifies the pixel-level segmentation accuracy by assessing the overlap between predicted and ground truth segmentation masks.

To further assess the model’s generalization ability, we compared its segmentation performance on wheat cultivars from the training and validation sets with those not included during training. This evaluation provides insights into the model’s robustness and adaptability to unseen cultivars, which is crucial for practical applications in diverse wheat populations.

## 3 Results

### 3.1 SpikePheno delivers accurate performance in wheat spike phenotyping

SpikePheno trained a spike segmentation model based on the ResNet50-UNet network to segment wheat spikes and stems from the background. The model achieved an mIoU of 0.950 and an mPA of 0.960 for seen cultivars, and 0.948 mIoU and 0.965 mPA for unseen cultivars (Table 2). These results demonstrate the model’s high accuracy and generalization ability, indicating that it performs slightly better on seen cultivars but maintains high reliability on unseen cultivars.

**Table 1.** Overview of traits derived for wheat spike and spikelet.

| Features | Abbreviation | Description |
| --- | --- | --- |
| Spike length | SL | Length of the main axis of the spike |
| Spike area | SA | Area of the spike (excluding awns) |
| Spike roundness | SR | Roundness of the spike (excluding awns) |
| Spikelet gap | SG | Average gap between adjacent spikelets of the spike |
| Spike width | SW | Average width of the spike |
| Length of fertile spikelet | LFS | Average length of fertile spikelets |
| Area of fertile spikelets | AFS | Average area of fertile spikelets |
| Roundness of fertile spikelets | RFS | Average roundness of fertile spikelets |
| Fertile spikelet angle | FSA | Average angle of fertile spikelets |
| Spikelet number per spike | SNS | Total spikelets count per spike |
| Infertile spikelet number | ISN | Number of infertile spikelets per spike |
| Fertile spikelet number | FSN | Number of fertile spikelets per spike |

**Table 2.**
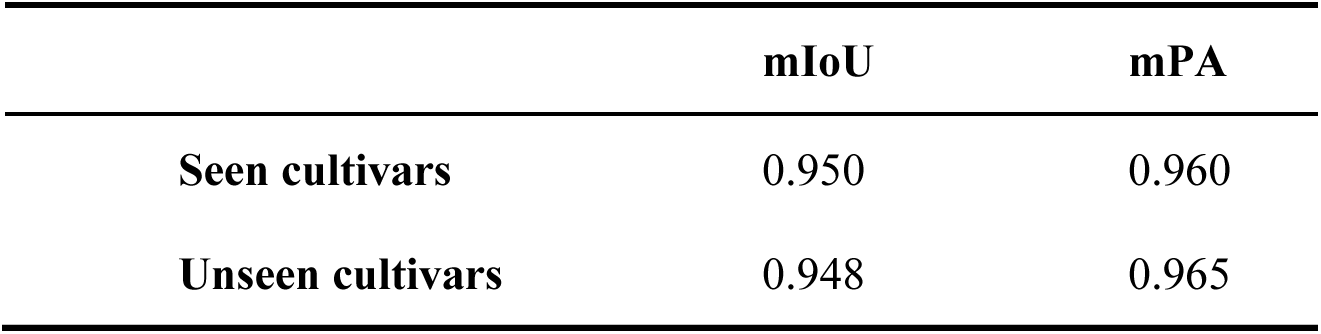
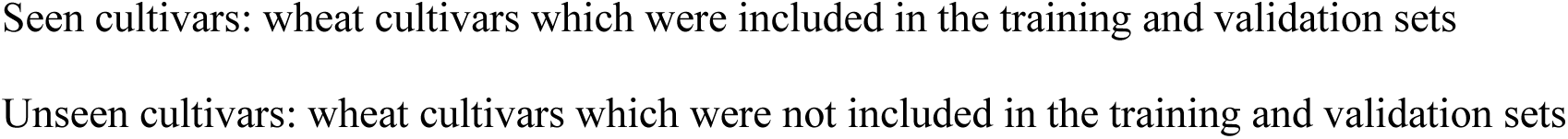
Performance of spike segmentation between seen and unseen wheat cultivars.

|  | mIoU | mPA |
| --- | --- | --- |
| Seen cultivars | 0.950 | 0.960 |
| Unseen cultivars | 0.948 | 0.965 |

For spikelet identification, we evaluated the performance of three different models—Mask R-CNN, PointRend [28], and Yolov8x-seg—on both seen and unseen wheat cultivars (Table 3). For unseen cultivars, Mask R-CNN achieved an mAP50box of 0.972 for detection and an mAP50seg of 0.973 for segmentation, compared to 0.956 mAP50box and 0.946 mAP50seg for seen cultivars. Similarly, PointRend reached an mAP50box of 0.979 and an mAP50seg of 0.979 for unseen cultivars, which is an improvement over the 0.948 mAP50box and 0.943 mAP50seg observed for seen cultivars. Yolov8x-seg performed exceptionally well on unseen cultivars, with an mAP50box of 0.986 and an mAP50seg of 0.986, noticeably higher than the 0.988 mAP50box and 0.984 mAP50seg for seen cultivars. These results highlight the models’ excellent generalization capabilities on wheat cultivars not directly trained on, and as a result, we have chosen Yolov8x-seg as our model for SpikePheno.

**Table 3.** Spikelets identification performance of different models between seen and unseen wheat cultivars.

|  | Seen cultivars |  | Unseen cultivars |  |
| --- | --- | --- | --- | --- |
|  | mAP50<sub>box</sub> | mAP50<sub>seg</sub> | mAP50<sub>box</sub> | mAP50<sub>seg</sub> |
| **Mask R-CNN** | 0.956 | 0.946 | 0.972 | 0.973 |
| **PointRend** | 0.948 | 0.943 | 0.979 | 0.979 |
| **Yolov8x-seg** | **0.988** | **0.984** | **0.986** | **0.986** |

Seen cultivars: wheat cultivars which were included in the training and validation sets Unseen cultivars: wheat cultivars which were not included in the training and validation sets

Seen cultivars: wheat cultivars which were included in the training and validation sets Unseen cultivars: wheat cultivars which were not included in the training and validation sets To further validate the reliability of our model, SNS counted manually in all collected images of wheat spikes was compared with the number predicted by the model (Fig. 4). The results demonstrate a high degree of consistency between the predicted and actual measured values. Regression analysis yielded a coefficient of determination (R^2^) of 0.9923 and an RMSE of 0.1581. These evaluation metrics emphasize the accuracy and reliability of our model in the task of spikelet counting, confirming its effectiveness in practical applications.

**Fig. 4.**
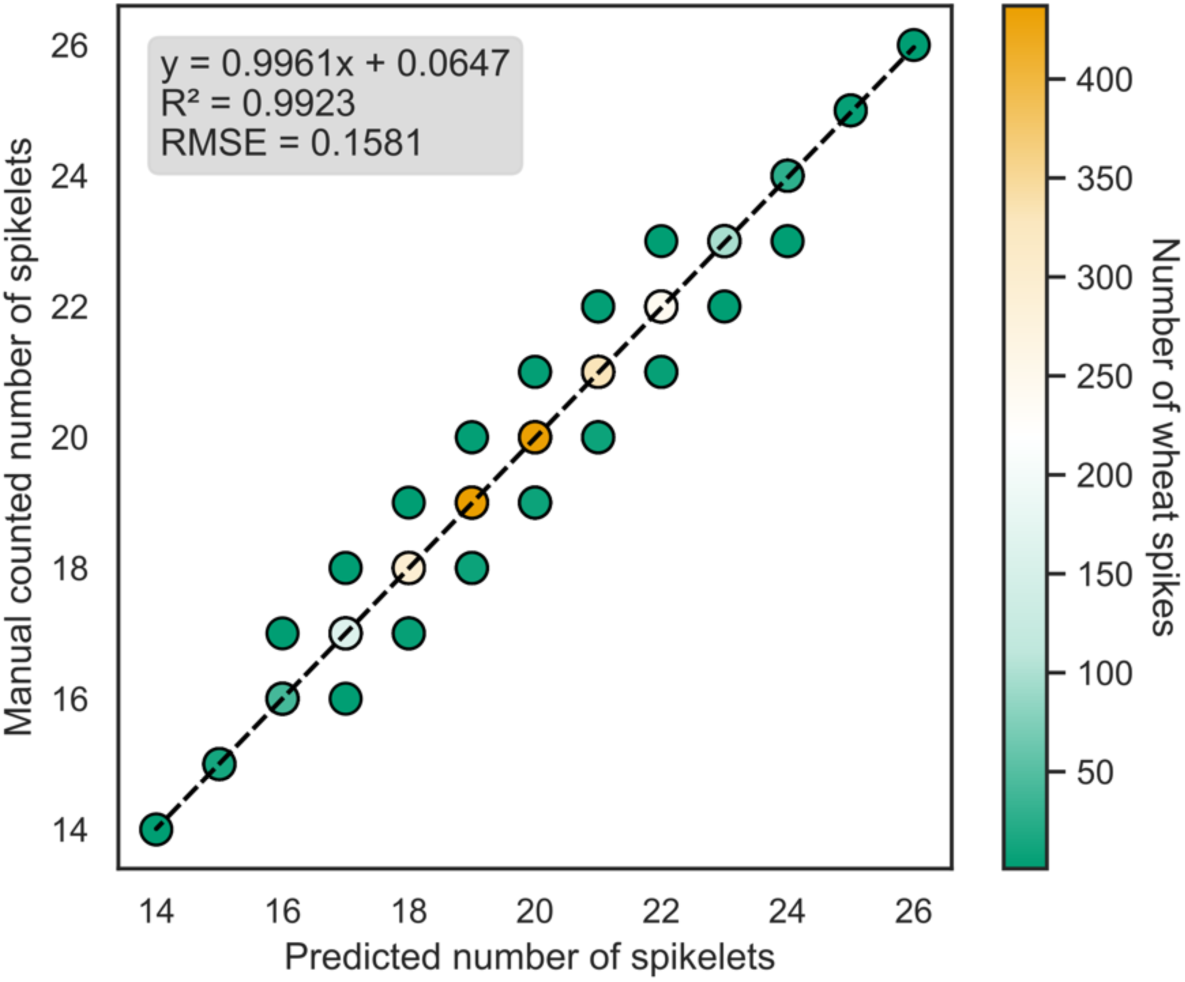
Comparison of manual counts and model prediction for spikelet number

### 3.2 Wheat spike morphological traits are correlated with TGW

We analyzed spike traits using a diverse set of 221 wheat accessions, which exhibited rich spike-type diversity. SpikePheno was used to extract 45 spike-related traits (Table S1). Of these, 12 key traits related to spike and spikelet architecture were selected for in-depth analysis, and their mean values are listed in Table S2. These traits exhibited extensive variation across the accessions, reflecting the phenotypic diversity present in the population (Fig. 5A).

**Fig. 5.**
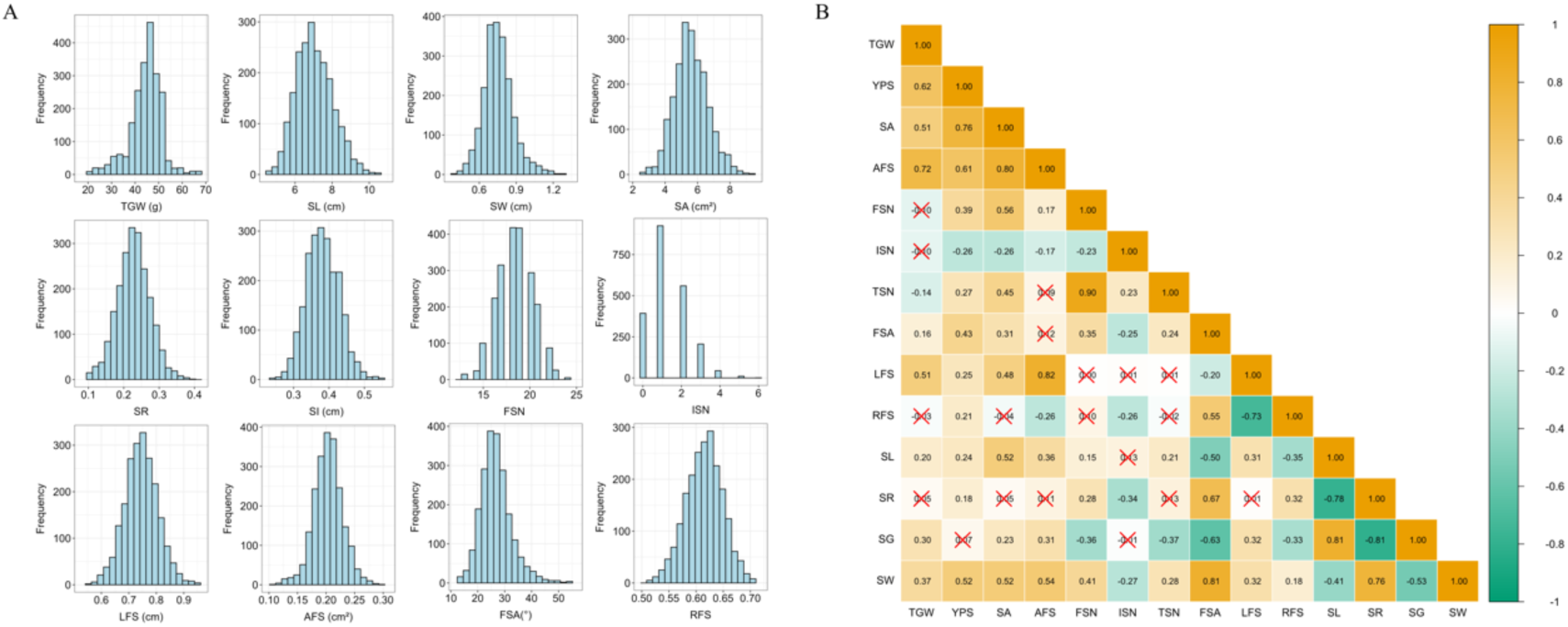
Distribution and correlation of wheat spike phenotypes. (A) Histograms of wheat spike phenotype distribution. (B) Correlation of different wheat spike phenotypes.

The high-resolution dissection of spike-related traits revealed significant correlations with yield-related traits (Fig. 5B). Among the extracted spike traits, SA showed a strong and positive correlation with both TGW (*r* = 0.51) and grain yield per spike (YPS, *r* = 0.76), indicating the high relevance of the extracted phenotypes to yield-related traits. SW also exhibited notable correlations with TGW (*r* = 0.37) and YPS (*r* = 0.52). In contrast, the traditional agronomic trait, SL, exhibited a much weaker correlation with TGW (*r* = 0.20) and YPS (*r* = 0.24). This emphasizes the superior utility of SA over SL for yield prediction. Similarly, for spikelet traits, the average area of fertile spikelet (AFS) showed a strong and positive correlation with TGW (*r* = 0.72) and YPS (*r* = 0.61).

Interestingly, AFS exhibited a stronger correlation with TGW than SA did. This is likely because TGW is a seed-level phenotype that is more closely related to spikelet-level traits like AFS. In contrast, SA showed a stronger correlation with YPS, which is a spike-level phenotype more directly tied to overall spike architecture. These findings suggest that both SA and AFS are critical for wheat yield, with SA influencing spike-level productivity and AFS directly affecting seed size and weight.

### 3.3 Large spikes and spikelets enhance wheat yield potential

We reduced the dimensionality of 45 phenotypes extracted from 221 cultivars of wheat spikelets into nine principal components (PCs) through principal component analysis (PCA), retaining 95% of the information to eliminate redundancy. Then, we used the complete linkage method to perform hierarchical clustering on the reduced data. Based on the clustering results, the spikes were categorized into 6 classes, each showcasing unique structural characteristics (Fig. 6A-B). In Class 1, the spikes are notably long with larger SAs, accompanied by a high number of medium-sized spikelets. These spikelets are characterized by narrower angles and larger gaps, resulting in a uniform yet open structure. In Class 2, the spikes are significantly shorter with smaller SAs and fewer spikelets. These spikelets are small in size and length but have moderately spaced gaps and angles, contributing to a compact and evenly distributed spike structure. In Class 3, the spikes tend to be longer with larger SAs and a moderate number of spikelets. The spikelets in this category are relatively larger and longer, with well-balanced angles and gaps, creating an evenly distributed spike structure. Class 4 spikes are shorter and have moderate SAs. The spikelets are fewer in number and moderate in size and length, but exhibit larger angles and smaller gaps, resulting in a more compact and uniform spike appearance. Class 5 spikes, while shorter, have the largest SAs. These spikelets are notably wide, with greater lengths, larger angles, and closer spacing, giving the spike a robust and tightly packed structure. Finally, Class 6 spikes are shorter and smaller, with fewer spikelets that display distinct top-to-bottom differences. The spikelets at the top are larger and longer, with wider angles and smaller gaps, creating a dense, clustered structure. In contrast, the bottom spikelets are smaller and shorter, with narrower angles and larger gaps, leading to a sparser distribution at the base of the spike.

**Fig. 6.**
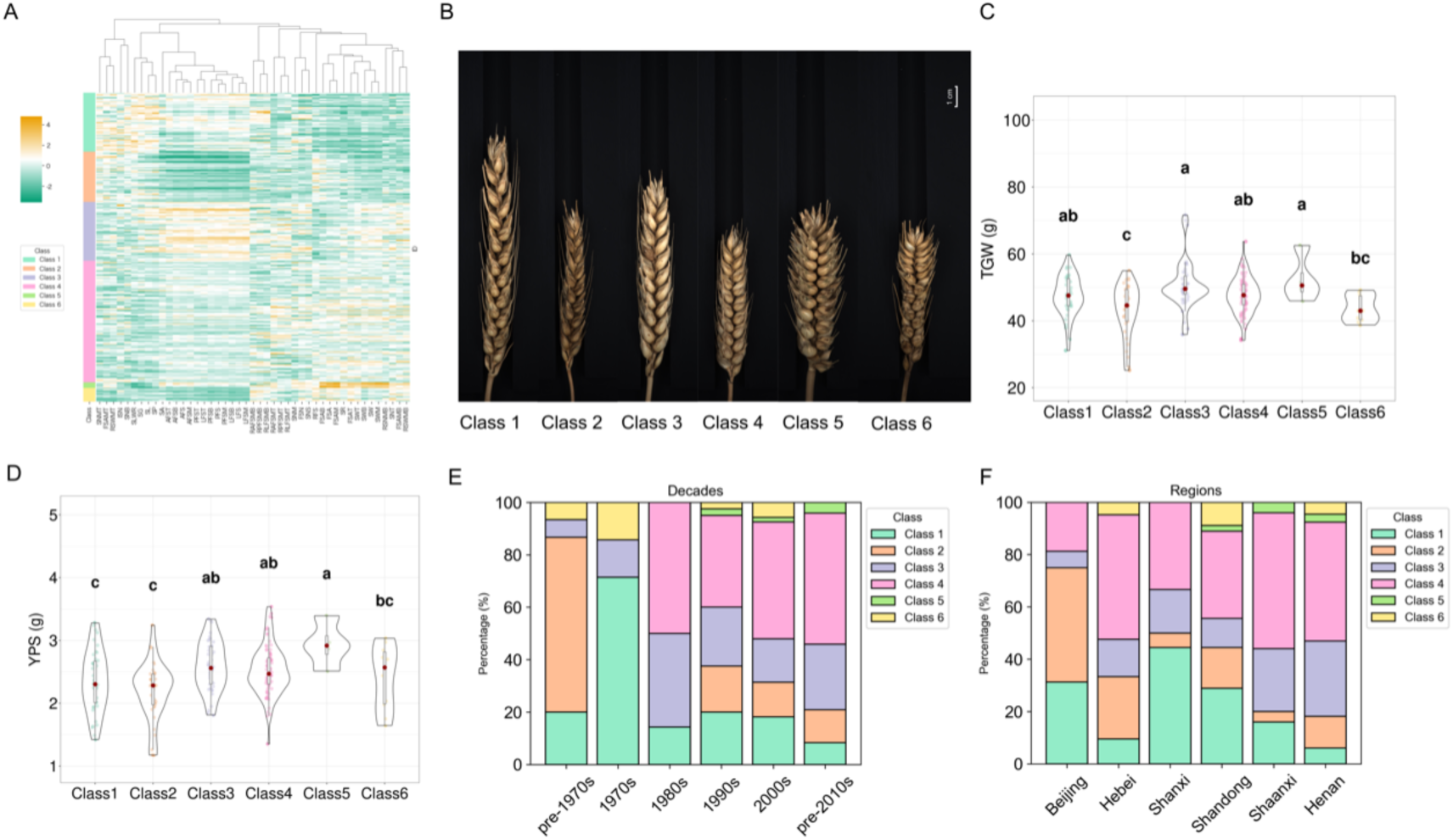
Analysis of spike classification and its temporal and regional differences in 221 wheat accessions. (A) Clustering heatmap of wheat spike phenotypes. (B) Representative spikes of the six classes of wheat spikes. (C) Thousand grain weight (TGW) of different classes of wheat spikes. (D) Yield per spike (YPS) of different classes of wheat spikes. (E) Distribution of classes across different decades. (F) Distribution of classes across different regions.

To assess the differences in yield potential among different spike categories, we conducted an analysis of variance (ANOVA) to preliminarily explore the statistical significance of TGW and YPS among classes. The results showed highly significant differences in TGW (*F* = 3.10, *p* < 0.01) and YPS (*F* = 5.37, *p* < 0.01) among these classes, indicating that different spike classes significantly influence TGW and YPS. To further determine the differences in TGW and YPS among these classes, we used the Least Significant Difference (LSD) test for multiple comparisons (Fig. 6C). For TGW, Classes 3 and 5 exhibited the highest values, with no statistically significant difference between them. Class 2 had the lowest TGW, which was significantly lower than that of all other classes except Class 6. Classes 1 and 4 showed intermediate TGW values, which were significantly higher than that of Class 2 but did not differ significantly from those of Classes 3, 5, and 6. Similarly, Class 6 had a lower TGW than Classes 3 and 5 but was not significantly different from other classes. For YPS, Classes 3, 4, and 5 exhibited the highest values. Classes 1 and 2 had the lowest YPS, which were significantly lower than those of all other classes except Class 6. Class 6 had an intermediate YPS, which was not significantly different from other classes except Class 5 (Fig. 6D).

The distribution of different classes has shifted significantly over time (Fig. 6E). Before the 1970s, Classes 1 and 2 were predominant, collectively accounting for most of the total percentage. However, starting from the 1980s, a notable increase in Classes 3 and 4 was observed, indicating a shift in dominance. Class 5, which was entirely absent before the 1980s, began to emerge during this period and remained low frequency afterward.

The Huang-Huai wheat region is one of the major wheat-producing areas in China. We analyzed the regional distribution of different wheat spike classes across the provinces of Beijing, Hebei, Shanxi, Shandong, Shaanxi, and Henan within this region. The distribution of these classes varied significantly across geographical regions (Fig. 6F). Class 4 was the dominant category in all regions, while Class 2 remain more prevalent in the northern provinces, such as Beijing and Hebei, but its proportions decrease moving southward. In contrast, Classes 3 and 4 gradually increase in proportion from north to south, with Class 3 being more prominent in Henan.

### 3.4 Long-term breeding trends: favoring large-spiked wheat cultivars

The morphological traits of wheat spikes and spikelets have changed over time (Fig. 7). TGW and YPS showed significant improvement before 1980s and remained stable afterwards. SA and SW increased gradually, with the lowest values before the 1970s and a noticeable rise from the 1980s onward, remaining high in recent decades. SL was higher in the 1970s compared to other decades, likely due to the larger proportion of Class 1 spikes (Fig. 6E), which are notably long with a larger SA and a high number of spikelets. After the 1980s, the increasing proportion of Classes 3 and 4 (Fig. 6E), which have moderate to large spikes and balanced structures, led to a more stable SL. AFS was highest in the 1980s and remained stable thereafter. The average length of fertile spikelets (LFS) and AFS were both at their lowest before the 1970s, likely due to the higher proportion of Class 2 (Fig. 6E), which was more prevalent during that period. Each fertile spikelet angle (FSA) remained low before the 1980s and showed a slight upward trend after the 1980s. This trend aligns with the increasing number of Classes 3 and 4 spikes and decreasing of Class 1 spikes, showing a similar pattern from the 1990s to the present. Therefore, the breeding of wheat has favored large-spike cultivars, with SA and key structural traits stabilizing at higher values. The dominance of these larger spike types in recent cultivars suggests a shift toward optimizing spike morphology for improved yield and adaptability.

**Fig. 7.**
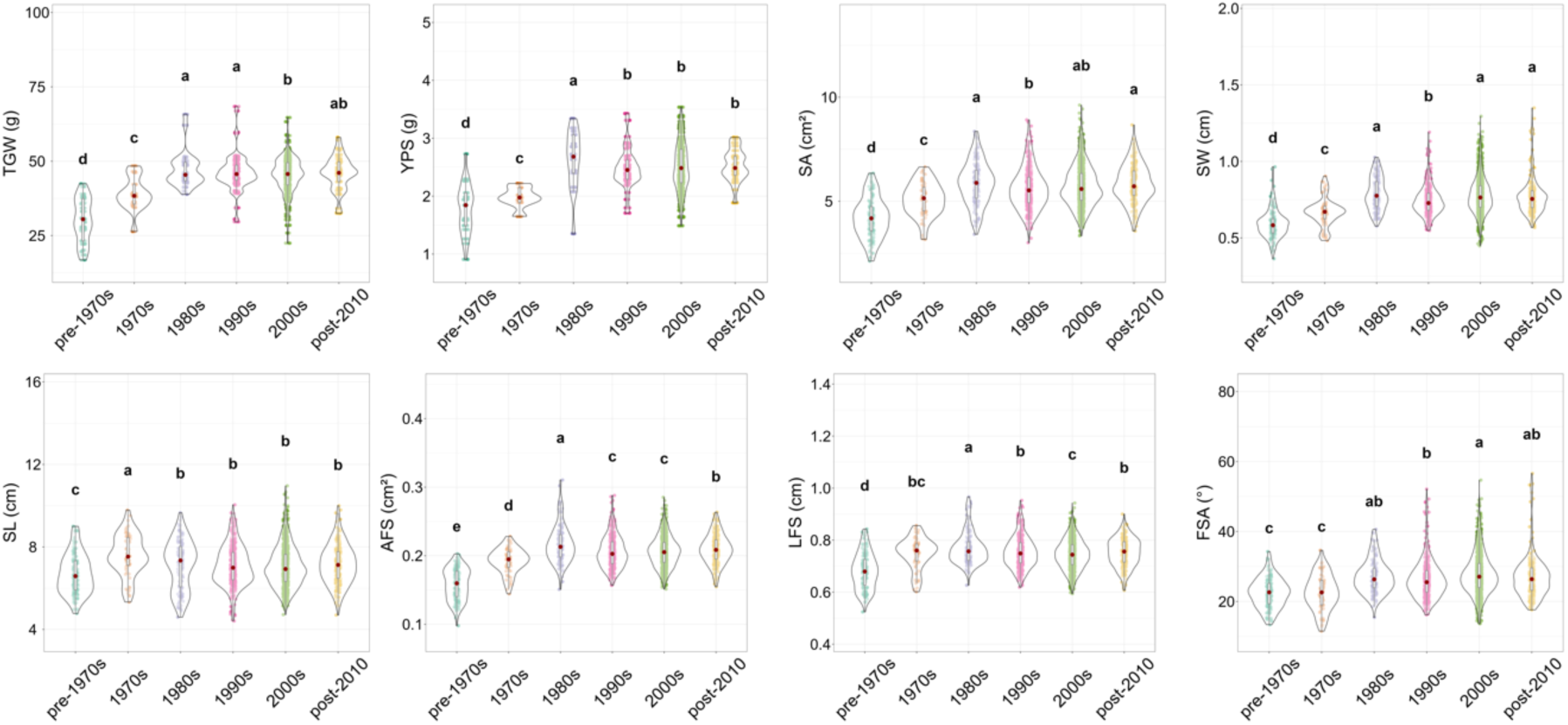
Differences in wheat spike phenotypes among decades.

### 3.4 Regional variation in wheat spike morphology across the Huang-Huai region, China

The morphological traits of wheat spikes and spikelets exhibited regional differences across the Huang-Huai wheat-production area of China, with notable variation among provinces (Fig. 8). SA and SW showed significant regional variation, with the highest values observed in Henan, a region in the southern part of the Huang-Huai wheat area which has the highest TGW and YPS. The lowest values were found in Beijing and Hebei, situated in the northernmost part of the region. The central provinces (Shanxi, Shandong, and Shaanxi) exhibited intermediate values, forming a transition between the extremes. SL, however, did not follow a clear geographical pattern, with some central provinces like wheats of Shanxi and Shandong displaying longer spikes, and wheats of Hebei and Henan had short SL.

**Fig. 8.**
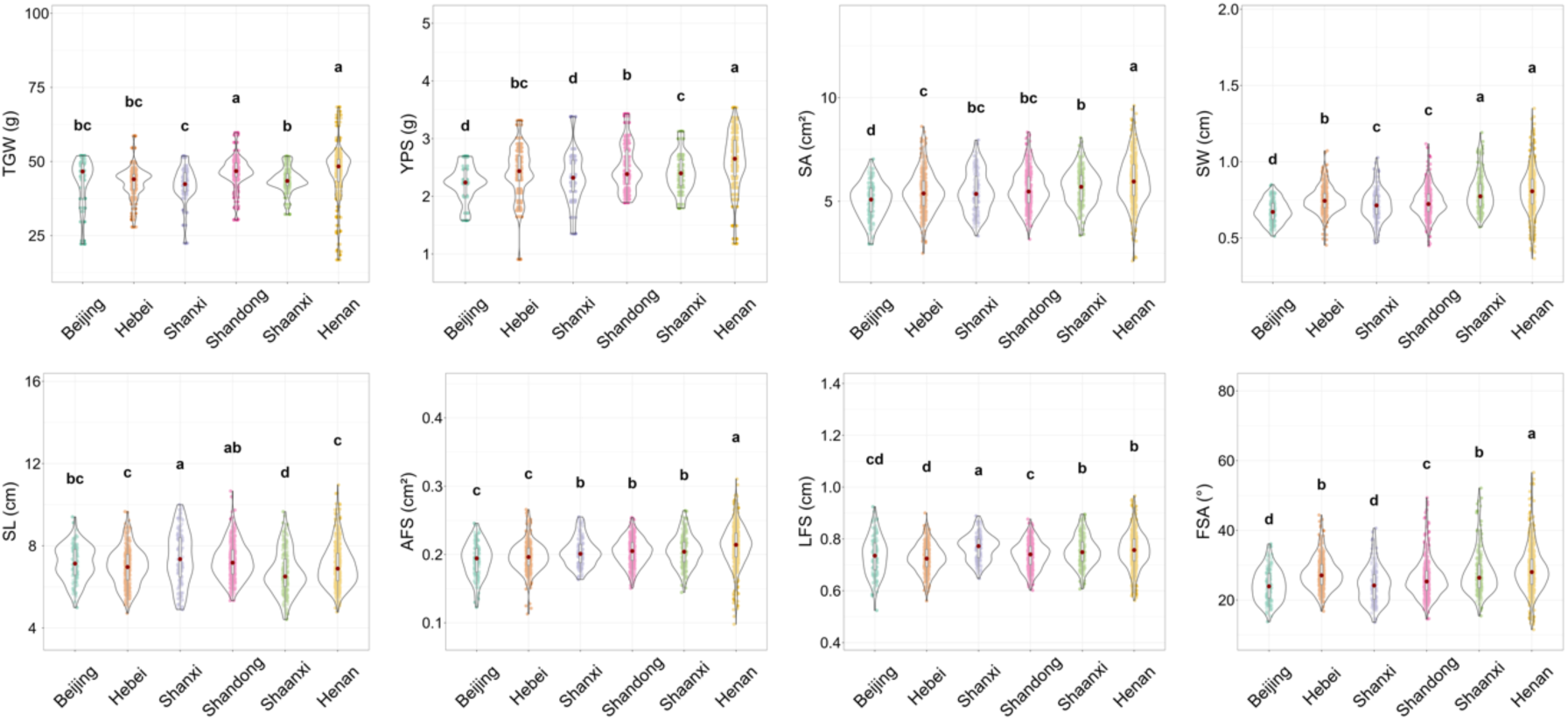
Differences in wheat spike phenotypes among regions.

Spikelet morphological characteristics also exhibited distinct regional variation (Fig. 8). AFS was highest in Henan and lowest in Beijing and Hebei, indicating a tendency for larger spikelets in the southernmost regions. LFS did not exhibit a clear regional distribution pattern, with variations observed across different provinces rather than a consistent north-south trend. The FSA was another key differentiator. The FSA of wheats in Henan was markedly greater than those in other provinces.

In the southern part of the Huang-Huai wheat region, Henan had a higher proportion of larger spikes and larger spikelets, whereas in the northernmost regions, Beijing and Hebei had a higher proportion of smaller spikes. This variation was closely linked to the distribution of spike classes. Hebei and Beijing contained a higher proportion of Class 2 spikes, which were characterized by smaller spikes with fewer and more compact spikelets. In contrast, Henan had a relatively higher proportion of Class 3 spikes, which were larger and had a more balanced structure (Fig. 6F). These differences in class composition contributed to the observed regional variation in spike and spikelet morphology.

## 4 Discussion

Wheat is an important global food crop, and its spike structure significantly impacts yield, pest and disease resistance, and stress tolerance [7, 29, 30]. Spikelets, as a crucial part of the wheat spike, significantly influence yield, and their shape and distribution on the spike also affect photosynthetic efficiency and grain filling [31]. We developed a wheat spike analysis pipeline based on image recognition that can precisely measure the spike morphological traits. We combined the ResNet50-UNet network and the YOLOv8x-seg network for accurate recognition of spikes and spikelets. Our approach achieved high accuracy in segmenting wheat spikes and stems, with mIoU of 0.950 and mPA of 0.966 for seen cultivars, and 0.948 mIoU and 0.965 mPA for unseen cultivars, demonstrating strong generalization ability (Table 2). Compared to existing methods, our approach can accurately distinguish between fertile and infertile spikelets using YOLOv8x-seg, which achieved an mAP50box of 0.986 and an mAP50seg of 0.986 on unseen cultivars (Table 3). Furthermore, our method extracts the main axis of the spike to accurately measure SL, ensuring a comprehensive and refined phenotypic analysis of spike architecture.

The comprehensive measurement and analysis of wheat spike morphology provide an essential foundation for understanding the mechanisms of yield formation [7]. Traditionally, it has been believed that longer spikes can offer more space and improved light conditions for grain development, thereby enhancing YPS. However, our correlation analysis results (Fig. 5) indicate SA and SW exhibited higher correlations (*r* = 0.76 and 0.68, respectively), whereas SL showed a correlation with YPS (*r* =0.42). Moreover, we found that the spikelet angle was also associated with YPS (*r* = 0.43), suggesting that a larger spikelet opening provides additional space for grain filling. Regarding TGW, our analysis revealed that AFS correlated more closely with TGW (*r* = 0.72) than does the overall SA (*r* = 0.51), implying that spikelet area—being directly related to individual grain development—better reflects variations in TGW. These findings underscore the importance of fine-scale spike traits—such as spikelet angle and spikelet area—in determining both YPS and TGW. Liu, Yu (30) also highlighted the value of detailed phenotypic analysis in understanding spike morphology, using 16 carefully manually measured traits to comprehensively describe the spike architecture. Their approach, which included detailed measurements such as spike weight, grain weight per spikelet, chaff weight per spike, the number of degenerated spikelets at both the top and the bottom of the spike, enhanced the ability to identify key loci associated with spike traits. This demonstrates that fine-scale phenotyping, whether through manual measurements or advanced imaging techniques, significantly improves the capacity to uncover the genetic basis of spike morphology and its impact on yield.

Nine PCs of 45 spike-related phenotypes enabled us to reduce the dimensionality of our data while retaining 95% of the original variability. Subsequent hierarchical clustering based these nine PCs identified six distinct clusters (Fig. 6A), each representing unique spike architectural patterns. When these clusters were analyzed in relation to the decades of cultivar release, a clear temporal trend emerged: older cultivars (pre-1970) were predominantly grouped in clusters characterized by longer yet more compact spikes, whereas cultivars released from the 1980s onward increasingly exhibited traits such as larger SA and greater SW, which are strongly correlated with enhanced YPS and TGW. This trend is consistent with the findings of Liu, Yu (30), who reported significant changes in spike morphological traits from 1900 to 2020, resulting in a fuller appearance due to increased fertile spikelet numbers. These changes reflect modern breeding efforts to optimize spike architecture for higher yield [32, 33]. Moreover, a regional analysis revealed that cultivars from Beijing and Hebei tended to possess spikes with higher density and compactness, while those from Henan province displayed wider spikes with larger spikelet areas (Fig. 8). This study is consistent with the findings of Zhang, Lu (34), who reported higher TGW in southern regions compared to northern regions. These findings indicate that both breeding history and regional adaptation have driven divergent trends in spike morphology, providing valuable insights for developing region-specific breeding strategies to optimize yield.

## 5 Conclusion

In conclusion, our study demonstrates that deep learning-based analysis of wheat spike morphology can provide detailed, high-throughput phenotypic data strongly associated with yield-related traits. By developing a two-stage imaging pipeline called SpikePheno that combines ResNet50-UNet for whole-spike semantic segmentation with YOLOv8x-seg for individual spikelet classification and segmentation, we achieved quantification of 45 morphological traits. Through this integrated approach, we demonstrate that fine-scale architectural features such as SL, SA, SW, and AFS are key determinants of YPS and TGW. The current pipeline can accurately identify infertile spikelets in the bottom part of the spike. However, its accuracy decreases when detecting infertile spikelets at the top of the wheat spikes, particularly where small spikelets are compressed. Improving imaging techniques is required to address this limitation in future research. Moreover, the principal component analysis and hierarchical clustering of spike traits revealed significant temporal and regional trends in spike morphology. Modern cultivars, particularly those from regions like Henan, tend to exhibit larger spike traits, such as increased spike area and spikelet area, which may be associated with higher yield potential. These findings enhance our understanding of the complex relationships between spike structure and yield formation and provide a valuable basis for precision breeding strategies aimed at developing high-yield wheat cultivars.

## Authors contributions

SZ, ZL and NJ designed and supervised the experiments. FS and QG performed the experiments. SZ collected the experimental data. FS developed the phenotyping pipeline. FS and QG analyzed the results. FS wrote the manuscript. SZ, ZYL, and NJ revised the manuscript. All authors read and approved the contents of this paper.

## Acknowledgments

This work was supported by the Biological Breeding-National Science and Technology Major Project (2023ZD04076), and the National Key Research and Development Program of China (2023YFF1000100).

## Data availability

The full implementation of the spikePheno pipeline is available in GitHub at URL: https://github.com/Jiang-Phenomics-Lab/spikePheno (the repository will be public after acceptance). For peer review, the code and model can be accessed here: https://1drv.ms/f/c/19a61e6557e1be9a/EsVe-Qj1OS5LsVIofq90-3IB1sE1J8fP35ZCLD55dqo-8A?e=ji2NZc.

